# ATX expression from GFAP^+^ cells is essential for embryonic development

**DOI:** 10.1101/2020.01.17.910141

**Authors:** Ioanna Sevastou, Ioanna Ninou, Vassilis Aidinis

## Abstract

Autotaxin (ATX) is secreted by various type of cells in health and disease and catalyzes the extracellular production of lysophosphatidic acid (LPA). In turn, LPA is a bioactive lysophospholipid promoting a wide array of cellular functions through its multiple G-protein coupled receptors, differentially expressed in almost all cell types. ATX expression has been shown necessary for embryonic development and has been suggested to participate in the pathogenesis of different chronic inflammatory diseases and cancer. Deregulated ATX and LPA levels have been reported in multiple sclerosis (MS) and its experimental model, experimental autoimmune encephalomyelitis (EAE). ATX genetic deletion from macrophages and microglia (CD11b^+^ cells) attenuated the severity of EAE, thus proposing a pathogenic role for the ATX/LPA axis in MS/EAE. In this report, increased ATX staining was localized to glial fibrillary acidic protein positive (GFAP^+^) cells, mostly astrocytes, in spinal cord sections from EAE mice at the peak of the disease. However, genetic deletion of ATX from GFAP^+^ cells resulted in embryonic lethality, suggesting a major role for ΑΤΧ expression from GFAP^+^ cells in embryonic development, that urges further dissection. Moreover, the re-expression of ATX from GFAP+ cells during the pathogenesis of EAE, reinforces the concept that ATX/LPA is a developmental program aberrantly reactivated upon chronic inflammation.

## Introduction

Autotaxin (ATX, ENPP2) is a secreted glycoprotein that catalyzes the extracellular synthesis of lysophosphatidic acid (LPA), a bioactive growth factor-like signaling lysophospholipid [1–3]. LPA signals through at least six receptors that exhibit widespread cell and tissue distribution and overlapping specificities [4]. LPA receptors couple with G-proteins leading to pleiotropic effects in a large variety of cell types, including cells of the central nervous system (CNS) [4].

ATX expression is necessary for embryonic development, participating through LPA production in the development of the vascular and neural systems. Ubiquitous genetic deletion of ATX results in embryonic lethality [5]. Moreover, ATX has been suggested to possess matricellular properties, modulating the physiology of oligodendrocytes as well as neuronal progenitors [6, 7]. In adult healthy life, the ATX γ isoform, that contains a 25 aa insert of unknown function, is highly expressed in the CNS [8, 9].

Increased ATX and LPA levels have been reported in different chronic inflammatory diseases and cancer [10]. In the CNS, ATX-dependent LPA signaling has been suggested to participate in initiation of neuropathic pain [4, 11], while increased ATX expression has been reported in neuroblastomas and glioblastomas [12]. Deregulated ATX and LPA levels have been reported, with some controversy, in the sera and cerebrospinal fluid (CSF) of patients with multiple sclerosis (MS) [13–17], as well as in mice with experimental autoimmune encephalomyelitis (EAE)[17, 18], suggesting a role for ATX/LPA in the pathogenesis of MS/EAE. Moreover, pharmacologic potent ATX inhibition has been reported to attenuate the development of EAE [19], further suggesting potential therapeutic benefits in targeting ATX in MS.

In the context of EAE, activated macrophages and microglia (F4/80^+^ CD11b^+^ cells) were shown to express ATX, and ATX genetic deletion from CD11b+ cells attenuated the severity of EAE, suggesting an overall pathogenic role for ATX in MS/EAE pathogenesis [18]. However, increased ATX levels have been detected in activated astrocytes following neurotrauma, completely absent in physiological conditions [20]. Astrocytes are the most abundant glial cells, responsible for physical and metabolic support of other neural cells. They participate in the regulation of immune responses in the CNS, and orchestrate tissue repair following injury [21]. Moreover, astrocytes are increasingly recognized as active players in myelination and MS pathogenesis [22, 23]. Therefore, in this report, we examined ATX expression from astrocytes during EAE development. Increased ATX staining was detected in activated astrocytes in the inflamed spinal cord upon EAE, suggesting yet another possible source of ATX expression in the inflamed CNS. However, genetic deletion of ATX from GFAP^+^ cells resulted to embryonic lethality, suggesting a major role of ATX expression from GFAP^+^ cells in embryonic development.

## Materials and Methods

### Mice

All animal work performed in this work was approved by the Institutional Animal Ethical Committee (IAEC) of BSRC Alexander Fleming and the Veterinary service and Fishery Department of the local governmental prefecture (#449 and #1369/3283/3204) in line with the ARRIVE guidelines. Mice were housed and bred at the BSRC Alexander Fleming animal facility at steady conditions of 20–22°C temperature, 55±5% humidity, and a 12-h light-dark cycle. Water and food were given *ad libitum*. The generation and genotyping protocols for *Enpp2*^f/f^ [5], and Tg*hGFAP-Cre* [24] genetically modified mice have been described previously. All mice were bred in the C57Bl6/J genetic background for over 10 generations. The health status of the mice was monitored at least once per day; no unexpected deaths were observed. Euthanasia was humanely performed in a CO_2_ chamber with gradual filling followed by exsanguination, at predetermined time-points. To identify embryonic lethality in TghGFAP-CreEnpp2^f/f^ mice, embryos from heterozygous *TghGFAP-CreEnpp2^+/−^* intercrosses were removed from the uterus and examined at different embryonic days post coitum.

### Experimental autoimmune encephalomyelitis (EAE)

EAE was induced in 10-12-week-old C57Bl6/J (H-2^b^) male mice following a widely used EAE protocol[25], as previously reported [26, 27]. Mice were subcutaneously immunized with 100 μg of 35-55 myelin oligodendrocyte glycoprotein (MOG35–55, MEVGWYRSPFSRVVHLYRNGK, GeneCust), emulsified in Freund’s Adjuvant supplemented with 1mg of heat-inactivated *Mycobacterium tuberculosis* H37RA (Difco Laboratories) to the side flanks. In addition, mice received two intraperitoneal injections of 100 ng pertussis toxin at the time of immunization and 48h later. Mice were weighed and monitored for clinical signs of EAE throughout the experiment. EAE symptoms were scored as follows: 0, no clinical disease; 1, tail weakness; 2, paraparesis (incomplete paralysis of 1 or 2 hind limbs); 3, paraplegia (complete paralysis of 1 or 2 hind limbs); 4, paraplegia with forelimb weakness or paralysis; 5, dead or moribund animal. At the day of sacrifice, blood plasma and spinal cord tissue were harvested and stored. All EAE experimental groups consisted of randomly assigned littermate male mice. All measures were taken to minimize animal suffering and distress; no invasive or painful techniques were performed requiring anesthetics or analgesics.

### Immunofluorescence

Mouse spinal cords were flushed out of the spinal canal, embedded in OCT and cryopreserved at −80°C. For immunofluorescence, 7 μm sections were sliced transversely onto super frost glass slides. Sections were fixed in 4% paraformaldehyde for 20 min at room temperature. Sections were then permeabilized with 0.2% Triton-X for 5 min for intracellular antigen detection wherever it was necessary. Non-specific antigen sites were blocked with blocking solution (Zytomed) for 5 min, followed by incubation with rabbit anti-mouse ATX (1:500, Cayman and/or Sigma) or rabbit IgG isotype control or mouse anti-glial fibrillary acidic protein (GFAP; clone G-A-5) conjugated with Cy3 (sigma, 1:2000) antibodies in 2% BSA at 4°C overnight. All washes were performed using PBS-Tween 0.05%. The following day anti-rabbit Alexa488 (Abcam, 1:1000) secondary antibody was applied to ATX stained sections for 1h at room temperature. All sections were counter-stained with DAPI (Fluoroshield with DAPI histology mounting medium, Sigma).

## Results and Discussion

### Increased ATX staining in spinal cord GFAP^+^ cells during EAE

Increased ATX levels have been reported in the spinal cords of mice developing EAE; some of this ATX increase was attributed to activated CD11b^+^ cell expression, mostly macrophages and microglia [18]. Here, we show that a robust ATX signal co-localizes with GFAP^+^ cells in spinal cord sections of relapsing as well as remitting EAE. As shown in Figure 1A, increased ATX expression was observed during EAE, peaking at the remission phase as previously reported [18]. Intense ATX staining was detected in astrocytes at the peak of the disease (GFAP^+^ cells; Fig. 1A), confirmed with confocal microscopy (Fig. 1B), suggesting that ATX is expressed from activated astrocytes during EAE. In line, increased ATX expression from activated astrocytes following neurotrauma has been previously reported [20]. However, some ATX staining could be attributed to soluble ATX bound to integrins or other transmembrane or membrane associated molecules [28–30].

**Figure 1.**
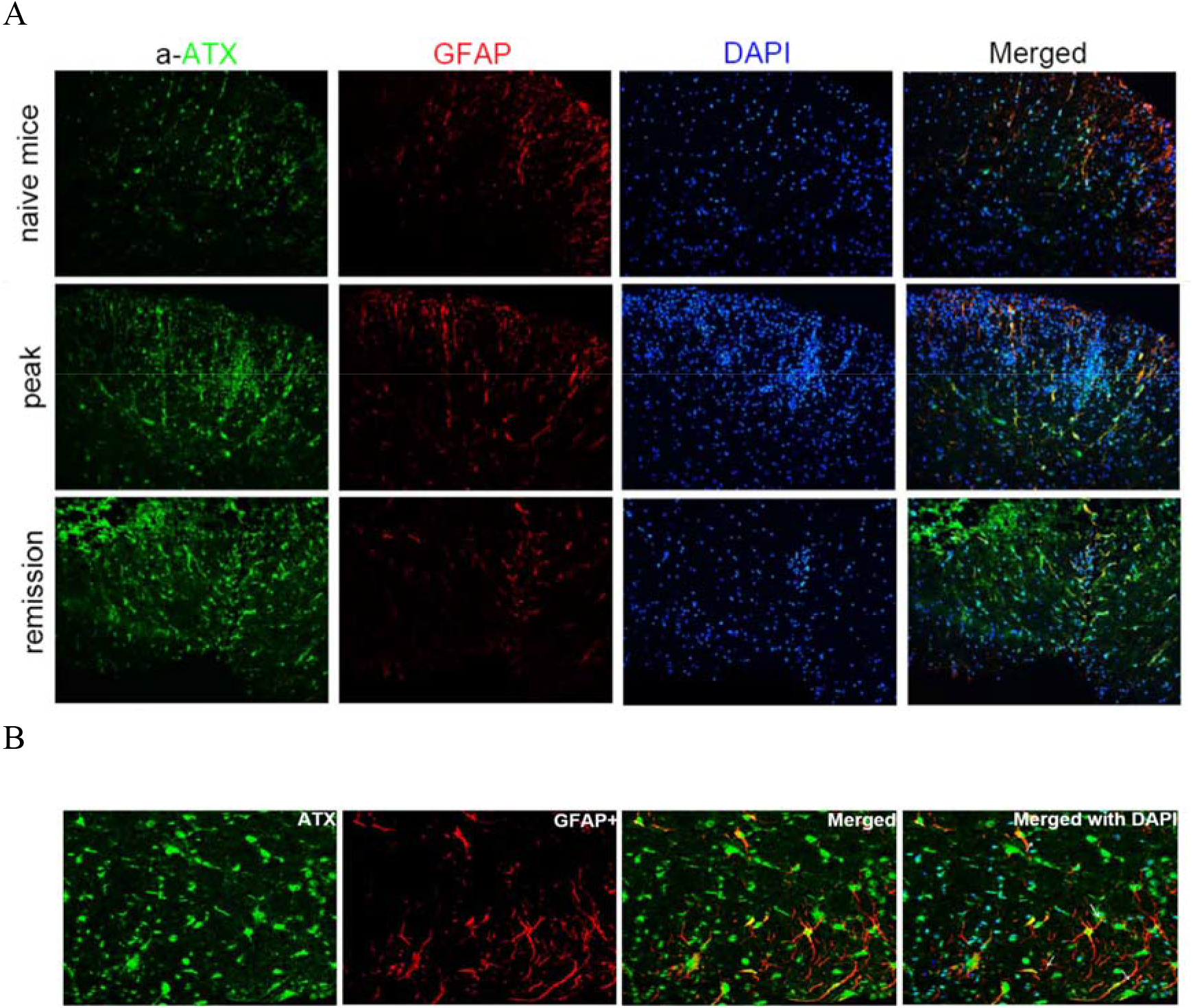
Increased spinal cord ATX staining upon EAE, partly localizing with astrocytes (GFAP^+^).

Astrocytic ATX expression could have different roles in EAE pathogenesis via the production of LPA. As the current dogma/hypothesis suggests, LPA forms and acts locally while different effects of LPA in astrocyte physiology have been suggested via well-established signal transduction pathways [4]. In brief, LPA can induce several of the hallmarks of reactive astrogliosis such as cytoskeletal re-organization [31–33], proliferation [34] and axonal outgrowth [35, 36]. Moreover, LPA has been reported to modulate astrocytic glucose uptake [37] and glycogenolysis [38], suggesting that LPA is involved in metabolic regulation of astrocytes and hence to the overall CNS energy metabolism, due to the central role of astrocytes in it. Other possible secondary, indirect effects of astrocytic ATX/LPA include neuronal differentiation [39] and possibly synaptic transmission [40] that require further exploration.

### ATX genetic deletion from GFAP^+^ cells results in embryonic lethality

In order to examine the role of ATX expression from reactive astrocytes in MS pathogenesis, ATX was genetically deleted from astrocytes by mating the conditional knockout mouse for ATX (*Enpp*2^f/f^)[5] with a transgenic mouse line expressing the Cre recombinase under the control of the hGFAP promoter (Tg*hGFAP-Cre*)[24], directing cre expression to astrocytes and GFAP^+^ neural stem cells. However, genetic deletion of ATX from GFAP^+^ cells resulted in embryonic lethality (Table 1, Fig. 2), irrespective of the origin (paternal or maternal) of the Cre transgene. It is worth noting that the very few *hGFAPCre^+/−^Enpp2^fl/fl^* mice that were born had not recombined the *Ennp2* gene, unlike the malformed and growth retarded embryos. Therefore, ATX expression from GFAP+ cells is necessary for embryonic development, extending previous studies indicating aberrant neural tube formation in ATX deficient embryos [5], whereas neural tube patterning is closely linked with specification and functions of astrocytes [21].

**Table 1:**
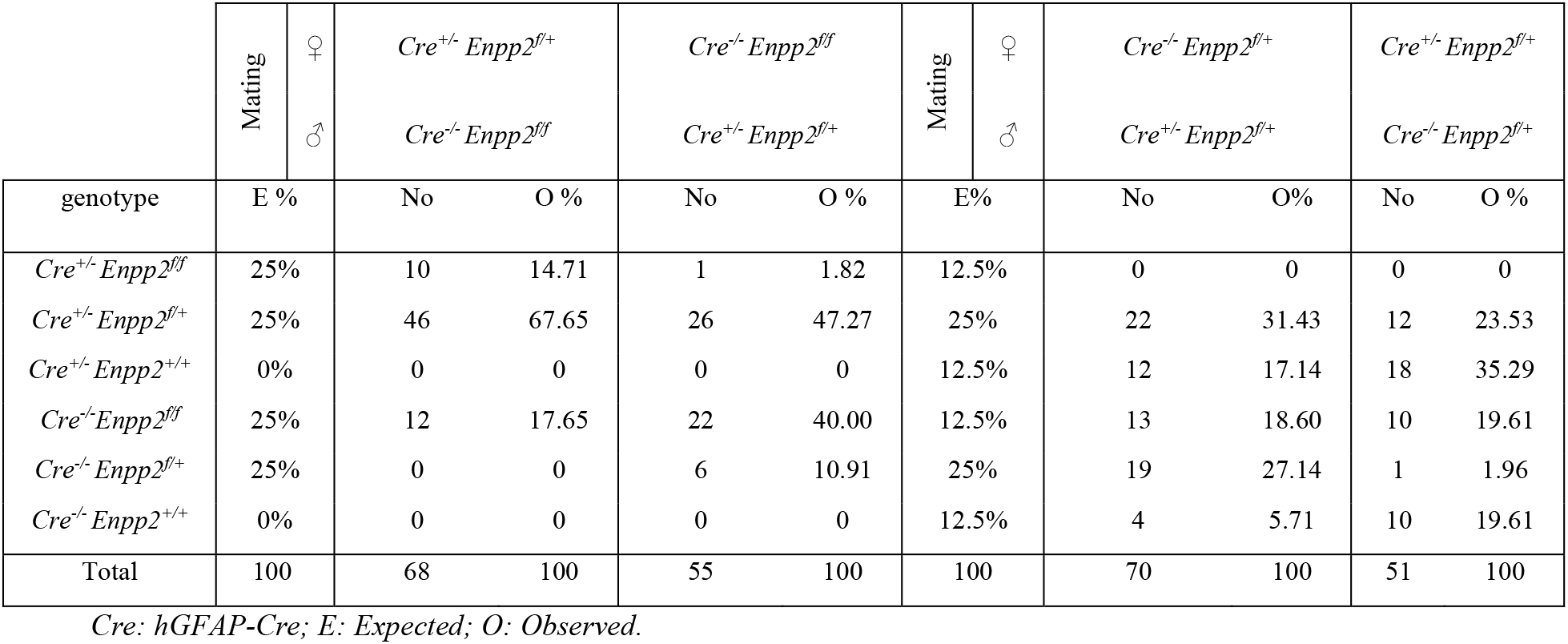
Genetic deletion of ATX from GFAP+ cells results in embryonic lethality. The expected (E) and the obtained (O) genotypes from the matings of the conditional knockout mouse for ATX (Enpp2^n/n^) with transgenic TghGFAP-Cre mice.

**Figure 2.**
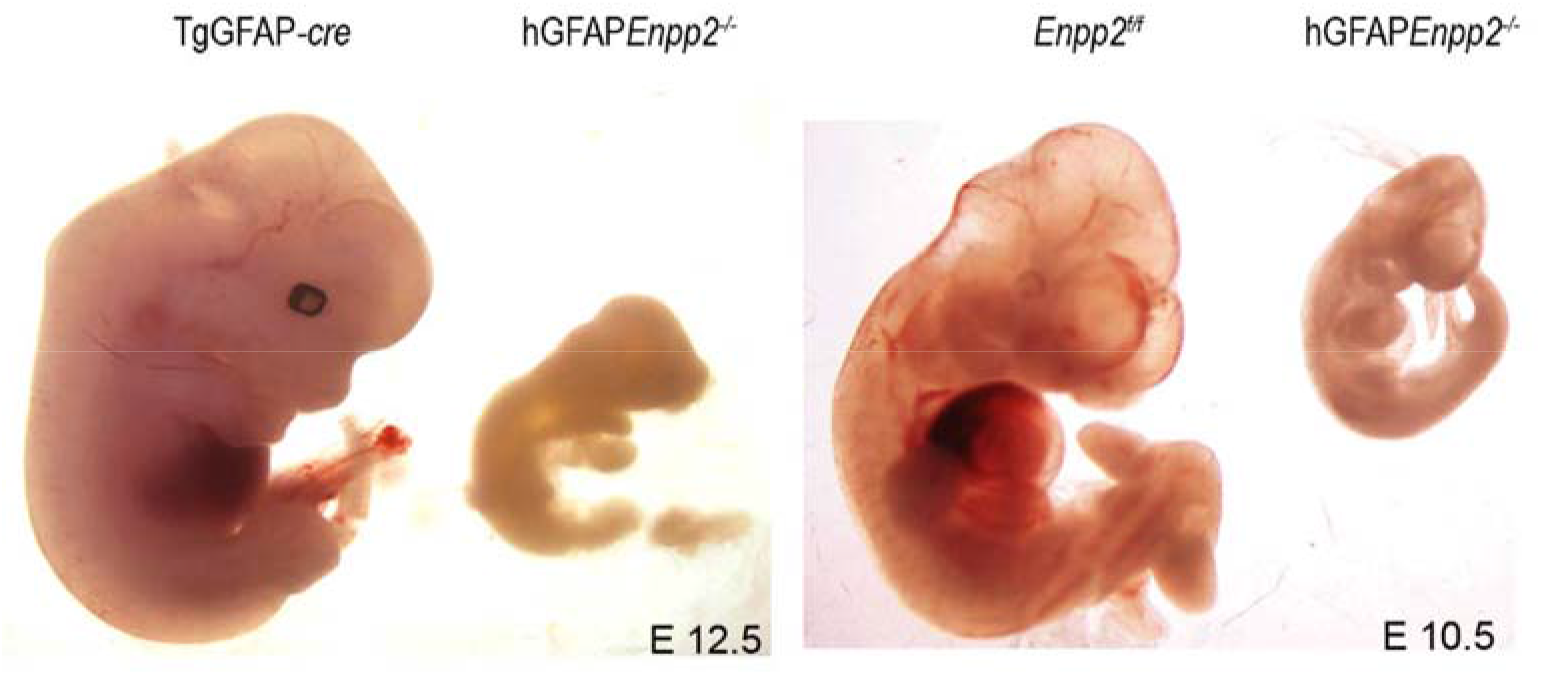
Genetic deletion of ATX from GFAP+ cells results to embryonic lethality.

The results presented here indicate that mature astrocytes produce ATX, as observed by cell staining following EAE induction (Figure 1). Additionally, ATX produced from GFAP expressing cells during embryonic development is necessary for the normal development of the embryo. This does not come as a surprise, given that in ATX deficient embryos the neural tube does not close, leading to embryonic lethality [5]. This implicates GFAP expressing cells as the primary sources of ATX in the early developing embryo. It has been shown that astrocytes start populating the developing brain at E16, with the majority of astrocytes being produced at the three first postnatal weeks of development [41]. We thus speculate that ATX is produced by GFAP^+^ progenitor cells that are present in the embryo at earlier gestational timepoints. GFAP^+^ progenitor cells in the mouse embryo are thought to exist as multifunctional cells called radial glia that can produce neurons [42] and astrocytes at a later stage [43]. Cre recombination of the GFAP promoter has first been detected as early as at E9.5 in the neuroepithelium and at E10.5 in a population of cells in the trigeminal ganglia [44].

ATX has been shown to be expressed at E8.5 of mouse development in the anterior folds of the neural tube and at the most posterior region of the midbrain but at E9.0-E9.5, ATX expression was strongly reduced in that brain region. At E10.5, prominent ATX expression was found at the floor plate of the neural tube [45]. This embryonic ATX expression may co-localise with early GFAP+ expressing cells of the neuroepithelium, resulting in the embryonic lethality observed in the mouse embryos expressing ATX under the hGFAP promoter. Further studies will be required to unveil the role of ATX expression from GFAP+ cells in early embryonic development. Regarding adult mouse studies, inducible deletion of ATX from GFAP expressing cells in adulthood could help to further elucidate the involvement of ATX in EAE pathogenesis.

## References

1. Barbayianni, E., et al., Autotaxin, a secreted lysophospholipase D, as a promising therapeutic target in chronic inflammation and cancer. Prog Lipid Res, 2015. 58: p. 76–96.

2. Ninou, I., C. Magkrioti, and V. Aidinis, Autotaxin in Pathophysiology and Pulmonary Fibrosis. Frontiers in Medicine, 2018.

3. Aikawa, S., et al., Lysophosphatidic acid as a lipid mediator with multiple biological actions. J Biochem, 2015. 157(2): p. 81–9.

4. Yung, Y.C., et al., Lysophosphatidic Acid signaling in the nervous system. Neuron, 2015. 85(4): p. 669–82.

5. Fotopoulou, S., et al., ATX expression and LPA signalling are vital for the development of the nervous system. Dev Biol, 2010. 339(2): p. 451–64.

6. Dennis, J., L. Nogaroli, and B. Fuss, Phosphodiesterase-Ialpha/autotaxin (PD-Ialpha/ATX): a multifunctional protein involved in central nervous system development and disease. J Neurosci Res, 2005. 82(6): p. 737–42.

7. Greenman, R., et al., Non-cell autonomous and non-catalytic activities of ATX in the developing brain. Front Neurosci, 2015. 9: p. 53.

8. Kanda, H., et al., Autotaxin, an ectoenzyme that produces lysophosphatidic acid, promotes the entry of lymphocytes into secondary lymphoid organs. Nat Immunol, 2008. 9(4): p. 415–23.

9. Giganti, A., et al., Murine and human autotaxin alpha, beta, and gamma isoforms: gene organization, tissue distribution, and biochemical characterization. J Biol Chem, 2008. 283(12): p. 7776–89.

10. Magkrioti, C., et al., Autotaxin and chronic inflammatory diseases. J Autoimmun, 2019. 104: p. 102327.

11. Inoue, M., et al., Initiation of neuropathic pain requires lysophosphatidic acid receptor signaling. Nat Med, 2004. 10(7): p. 712–8.

12. Yuelling, L.M. and B. Fuss, Autotaxin (ATX): A multi-functional and multi-modular protein possessing enzymatic lysoPLD activity and matricellular properties. Biochim Biophys Acta, 2008. 1781(9): p. 525–30.

13. Hammack, B.N., et al., Proteomic analysis of multiple sclerosis cerebrospinal fluid. Mult Scler, 2004. 10(3): p. 245–60.

14. Zahednasab, H., et al., Increased autotaxin activity in multiple sclerosis. J Neuroimmunol, 2014. 273(1-2): p. 120–3.

15. Jiang, D., et al., Elevated lysophosphatidic acid levels in the serum and cerebrospinal fluid in patients with multiple sclerosis: therapeutic response and clinical implication. Neurological Research, 2018: p. 1–5.

16. Balood, M., et al., Elevated serum levels of lysophosphatidic acid in patients with multiple sclerosis. Hum Immunol, 2014. 75(5): p. 411–3.

17. Schmitz, K., et al., Dysregulation of lysophosphatidic acids in multiple sclerosis and autoimmune encephalomyelitis. Acta Neuropathologica Communications, 2017. 5(1): p. 42.

18. Ninou, I., et al., Genetic deletion of Autotaxin from CD11b+ cells decreases the severity of experimental autoimmune encephalomyelitis. bioRxiv, 2019: p. 850149.

19. Thirunavukkarasu, K., et al., Pharmacological characterization of a potent inhibitor of autotaxin in animal models of inflammatory bowel disease and multiple sclerosis. J Pharmacol Exp Ther, 2016.

20. Savaskan, N.E., et al., Autotaxin (NPP-2) in the brain: cell type-specific expression and regulation during development and after neurotrauma. Cell Mol Life Sci, 2007. 64.

21. Molofsky, A.V., et al., Astrocytes and disease: a neurodevelopmental perspective. Genes Dev, 2012. 26(9): p. 891–907.

22. Kiray, H., et al., The multifaceted role of astrocytes in regulating myelination. Exp Neurol, 2016. 283(Pt B): p. 541–9.

23. Ponath, G., C. Park, and D. Pitt, The Role of Astrocytes in Multiple Sclerosis. Front Immunol, 2018. 9: p. 217.

24. Zhuo, L., et al., hGFAP-cre transgenic mice for manipulation of glial and neuronal function in vivo. Genesis, 2001. 31(2): p. 85–94.

25. Constantinescu, C.S., et al., Experimental autoimmune encephalomyelitis (EAE) as a model for multiple sclerosis (MS). Br J Pharmacol, 2011. 164(4): p. 1079–106.

26. Kassiotis, G. and G. Kollias, Uncoupling the proinflammatory from the immunosuppressive properties of tumor necrosis factor (TNF) at the p55 TNF receptor level: implications for pathogenesis and therapy of autoimmune demyelination. J Exp Med, 2001. 193(4): p. 427–34.

27. Tsakiri, N., et al., TNFR2 on non-haematopoietic cells is required for Foxp3+ Treg-cell function and disease suppression in EAE. Eur J Immunol, 2012. 42(2): p. 403–12.

28. Hausmann, J., et al., Structural basis of substrate discrimination and integrin binding by autotaxin. Nat Struct Mol Biol, 2011. 18(2): p. 198–204.

29. Leblanc, R., et al., Interaction of platelet-derived autotaxin with tumor integrin alphaVbeta3 controls metastasis of breast cancer cells to bone. Blood, 2014.

30. Fulkerson, Z., et al., Binding of autotaxin to integrins localizes lysophosphatidic acid production to platelets and mammalian cells. The Journal of biological chemistry, 2011. 286(40): p. 34654–63.

31. Guasch, R.M., et al., RhoA and lysophosphatidic acid are involved in the actin cytoskeleton reorganization of astrocytes exposed to ethanol. J Neurosci Res, 2003. 72(4): p. 487–502.

32. Manning, T.J., Jr., S.S. Rosenfeld, and H. Sontheimer, Lysophosphatidic acid stimulates actomyosin contraction in astrocytes. J Neurosci Res, 1998. 53(3): p. 343–52.

33. Ramakers, G.J. and W.H. Moolenaar, Regulation of astrocyte morphology by RhoA and lysophosphatidic acid. Exp Cell Res, 1998. 245(2): p. 252–62.

34. Shano, S., et al., Lysophosphatidic acid stimulates astrocyte proliferation through LPA1. Neurochem Int, 2008. 52(1-2): p. 216–20.

35. Spohr, T.C., et al., LPA-primed astrocytes induce axonal outgrowth of cortical progenitors by activating PKA signaling pathways and modulating extracellular matrix proteins. Front Cell Neurosci, 2014. 8: p. 296.

36. Spohr, T.C., et al., Astrocytes treated by lysophosphatidic acid induce axonal outgrowth of cortical progenitors through extracellular matrix protein and epidermal growth factor signaling pathway. J Neurochem, 2011. 119(1): p. 113–23.

37. Tomas, M., et al., Protective effects of lysophosphatidic acid (LPA) on chronic ethanol-induced injuries to the cytoskeleton and on glucose uptake in rat astrocytes. J Neurochem, 2003. 87(1): p. 220–9.

38. Choi, S.H., et al., Gintonin, a Ginseng-Derived Exogenous Lysophosphatidic Acid Receptor Ligand, Protects Astrocytes from Hypoxic and Re-oxygenation Stresses Through Stimulation of Astrocytic Glycogenolysis. Mol Neurobiol, 2018.

39. Spohr, T.C., et al., Lysophosphatidic acid receptor-dependent secondary effects via astrocytes promote neuronal differentiation. J Biol Chem, 2008. 283(12): p. 7470–9.

40. Thalman, C., et al., Astrocytic ATX fuels synaptic phospholipid signaling involved in psychiatric disorders. Molecular Psychiatry, 2018. 23(8): p. 1685–1686.

41. Ge, W.P., et al., Local generation of glia is a major astrocyte source in postnatal cortex. Nature, 2012. 484(7394): p. 376–80.

42. Malatesta, P., et al., Neuronal or glial progeny: regional differences in radial glia fate. Neuron, 2003. 37(5): p. 751–64.

43. Guo, Z., et al., Early postnatal GFAP-expressing cells produce multilineage progeny in cerebrum and astrocytes in cerebellum of adult mice. Brain Res, 2013. 1532: p. 14–20.

44. Andrae, J., et al., A 1.8kb GFAP-promoter fragment is active in specific regions of the embryonic CNS. Mech Dev, 2001. 107(1-2): p. 181–5.

45. Moolenaar, W.H., et al., Autotaxin in embryonic development. Biochimica et biophysica acta, 2012.

